# Cross-tissue comparison of telomere length and quality metrics of DNA among individuals aged 8 to 70 years

**DOI:** 10.1101/2023.08.19.553973

**Authors:** Sarah E. Wolf, Waylon J. Hastings, Qiaofeng Ye, Laura Etzel, Abner T. Apsley, Christopher Chiaro, Christine C. Heim, Thomas Heller, Jennie G. Noll, Hannah M.C. Schreier, Chad E. Shenk, Idan Shalev

## Abstract

Telomere length (TL) is an important biomarker of cellular aging, yet its links with health outcomes may be complicated by use of different tissues. We evaluated within- and between-individual variability in TL and quality metrics of DNA across five tissues using a cross-sectional dataset ranging from 8 to 70 years (N=197). DNA was extracted from all tissue cells using the Gentra Puregene DNA Extraction Kit. Absolute TL (aTL) in kilobase pairs was measured in buccal epithelial cells, saliva, dried blood spots (DBS), buffy coat, and peripheral blood mononuclear cells (PBMCs) using qPCR. aTL significantly shortened with age for all tissues except saliva and buffy coat, although buffy coat was available for a restricted age range (8 to 15 years). aTL did not significantly differ across blood-based tissues (DBS, buffy coat, PBMC), which had significantly longer aTL than buccal cells and saliva. Additionally, aTL was significantly correlated for the majority of tissue pairs, with partial Spearman’s correlations controlling for age and sex ranging from ⍴ = 0.18 to 0.51. We also measured quality metrics of DNA including integrity, purity, and quantity of extracted DNA from all tissues and explored whether controlling for DNA metrics improved predictions of aTL. We found significant tissue variation: DNA from blood-based tissues had high DNA integrity, more acceptable A260/280 and A260/230 values, and greater extracted DNA concentrations compared to buccal cells and saliva. Longer aTL was associated with lower DNA integrity, higher extracted DNA concentrations, and higher A260/230, particularly for saliva. Model comparisons suggested that incorporation of quality DNA metrics improves models of TL, although relevant metrics vary by tissue. These findings highlight the merits of using blood-based tissues and suggest that incorporation of quality DNA metrics as control variables in population-based studies can improve TL predictions, especially for more variable tissues like buccal and saliva.

## Introduction

Characterizing variation in telomere length (TL) and its links to human health outcomes is of interest across diverse scientific disciplines. Telomeres are ribonucleoprotein structures that maintain and protect the ends of chromosomes [1]. Telomeres shorten during cell division, resulting in age-related decreases in TL [2–4], occurring most rapidly early in life and continuing across the lifespan [5]. Variable TLs among same-aged individuals are thought to be the result of inherited genetic determinants of TL [6–8] and environmental exposures that accelerate TL loss [9–12]. Because short TL is linked to higher risk of age-related health outcomes [13–16] and early mortality [17–19], TL is frequently used as a biomarker of cellular aging in population studies [20, 21]. However, applications of TL to assess morbidity and mortality risk have produced inconsistent findings [19], leading to concerns about the utility of TL as a biomarker of aging [22, 23]. Importantly, inconsistencies in population research may be driven by key methodological differences in study design (e.g., tissue type, covariates selection, DNA extraction) [24–27].

TL has the potential to be an important biomarker of cellular aging in epidemiological and clinical research, yet establishing clear links with health outcomes are complicated by the use of different tissues across studies. Blood leukocytes, peripheral blood mononuclear cells (PBMCs), dried blood spots (DBS), saliva, and buccal epithelial cells are commonly used in population-based studies. Within an individual, TL may vary among these tissues due to factors such as cell composition, cell turnover rates, stem cell capacity to regenerate or differentiate, and dynamic regulation of TL by telomerase and other associated proteins [28–33]. Previous work has shown TL appears moderately to strongly correlated across tissues [0.53 < r < 0.93; 34, 35-37], although sex, behaviors (e.g., smoking), and telomere measurement assay may modulate these patterns [35, 36, 38]. Moreover, despite being correlated, there appear to be significant differences in measured TL across tissues [34–37, 39]. For example, Demanelis et al. [35] showed that tissue type accounted for 11.5-24.3% of variation in measured TL, which clustered by the developmental origin of each tissue. McLester-Davis et al. [38] demonstrated similar findings in a previous meta-analysis, observing stronger correlations among related tissues, e.g., blood-based tissues. Importantly, this meta-analysis also noted significantly lower correlations between tissues collected peripherally (e.g., buccal, PBMCs) and those collected surgically (e.g., bone marrow, spleen), highlighting the importance of tissue collection and processing procedures in cross-tissue concordance of TL measurements. In addition, previous work has also demonstrated significant differences in quality metrics of DNA across different tissues [40, 41], however it remains uncertain to what degree tissue-specific variation in the integrity, purity, and quantity of extracted DNA may influence the efficacy of TL assays and correlations among tissues. Given that tissue type is often a significant moderator of associations between TL and health outcomes [42, 43], it is vital that we better understand tissue diversity in TL.

Here, we quantified variation in absolute TL (aTL) across five tissues that are commonly used in population studies, namely buccal epithelial cells, saliva, DBS, buffy coat (i.e., leukocytes), and PBMCs. We evaluated within-and between-individual variation in aTL using a cross-sectional dataset of individuals ranging from 8 to 70 years of age. First, we quantified biological variation in aTL across tissues, age, sex, and race. We next evaluated whether tissues varied in the integrity, purity, and quantity of extracted DNA, which may influence the success and precision of telomere measurement assays. We subsequently assessed whether inclusion of information about DNA integrity, purity, and quantity improves model fits of aTL. Finally, we make recommendations on an optimal tissue type and quality control guidelines of extracted DNA for large population-based research.

## Materials and Methods

### Study Design and Sample Recruitment

Study participants were recruited from the Pennsylvania State University (PSU) community and surrounding areas, with some children recruited from other regions within Pennsylvania, as described in more detail below. This study and all protocols were approved by PSU’s Institutional Review Board.

### Adults

Adult participants were recruited via advertisements located on PSU’s University Park campus, community bulletins in State College and surrounding areas. Inclusion criteria for the study included: (a) ages 18-75, (b) no significant medical illness or immune disease (e.g., cancer, diabetes, or autoimmune disease), (c) current non-smoker, and (d) not pregnant or currently breastfeeding. Individuals were excluded if they self-reported a recent infection, illness, and/or use of antibiotics. To balance across ages and sex, eligibility became more restricted as sampling progressed. The maximum age was restricted to 75 years due to mortality selection [44] and the longer telomeres in exceptionally old individuals compared with controls with advancing age [45]. This study included 77 adult participants between 18 and 70 years old (**Table 1**).

**Table 1.** Demographic summary of participants, split by child and adult cohorts.

|  | Child (n = 120) | Adult (n = 77) |
| --- | --- | --- |
|  | Mean (SD) / Min-Max / N (%) |  |
| Age (years) | 11.95 (1.50) | 42.45 (15.70) |
| Age Range (years) | 8.6-15.08 | 18.28-70.01 |
| Sex |  |  |
| Female | 246 (51.5%) | 168 (54.5%) |
| Male | 232 (48.5%) | 140 (45.5%) |
| Race |  |  |
| White | 334 (69.9%) | 264 (86.8%) |
| Black | 52 (10.9%) | 8 (2.6%) |
| Other | 92 (19.2%) | 32 (10.5%) |

After obtaining informed consent, tissue samples and demographic information were collected from adult participants at PSU’s Clinical Research Center (CRC). First, participants completed a set of paper questionnaires to collect demographic and health-related information. Second, four tissue cells were collected, namely PBMCs, DBS, saliva, and buccal cells. Specifically, 20 mL of whole blood was collected in EDTA tubes via antecubital venipuncture by a trained phlebotomist. Approx. 200 µL of whole blood was applied to a Whatman 903 protein saver card, which we refer to as a dried blood spot (i.e., “DBS”), after which PBMCs were isolated through density-gradient centrifugation using Ficoll. Participants were also asked to provide 4 mL of saliva across two Oragene tubes (OGR-500, DNA Genotek), which upon completion, was mixed with the Oragene stabilizing buffer and sealed. Last, buccal cells were collected non-invasively using sanitary swabs (Isohelix SK1; 8 per individual), which were coated in cells by firmly scraping against the inside of the cheek several times in each direction. Collection order for all tissue types was uniform across participants. Participants were asked to refrain from eating or drinking anything other than water for one hour before arriving at the CRC. Tissue samples were then stored as follows: PBMCs were stored at −80°C in a solution buffer composed of phosphate buffered saline pH 7.2+EDTA (2mMol) + bovine serum albumin (0.5%) prior to extraction. DBS were stored in sealed Ziploc bags with desiccant packets at room temperature. Buccal swabs were placed in sealed Ziploc bags and stored at −80°C. Saliva samples were aliquoted into 4 cryovials and stored at −80°C.

### Children

Child participants were members of the Child Health Study (CHS), a large multidisciplinary study designed to provide prospective, longitudinal data on the health and development of children with and without a history of maltreatment investigations [for more details about the CHS see 46]. Children were recruited using the PA statewide Child Welfare Information System (CWIS) for having been investigated for substantiated maltreatment (i.e., defined according to PA state law, including sexual abuse, physical abuse and neglect) within the past year, and a demographically matched group of control children screened via CWIS to ensure no history of child welfare involvement. While the CHS study is recruiting 700 children, this investigation included the first 120 children enrolled between the ages of 8 to 15 years (**Table 1**).

Non-maltreating caregivers accompanied children to PSU’s University Park campus. After obtaining informed consent (caregiver) and assent (child), tissue samples and health/demographic data were collected from child participants. Four tissue cells were collected, namely buffy coat, DBS, saliva, and buccal cells. Specifically, 20 mL of whole blood was collected in EDTA tubes via antecubital venipuncture by a trained phlebotomist. Buffy coat was isolated using centrifugation to separate plasma followed by treatment with 0.5x red blood cell lysis buffer (Invitrogen). Using identical procedures to those described in adults, approx. 200 µL of whole blood was used to collect a DBS sample on a Whatman 903 protein saver card, and 2 mL of saliva (Oragene OGR-500, DNA Genotek) and 2 buccal cheek swabs (Isohelix SK1) were also taken per individual. Tissue samples were stored in the same conditions as adult samples, and buffy coat was stored at −80°C in a solution buffer composed of phosphate buffered saline pH 7.2+EDTA (2mMol) + bovine serum albumin (0.5%).

### Demographic measures

Chronological age, sex, and race were included as covariates because they are commonly associated with TL [2, 47–49]. Biological sex was determined via self-report. Race was coded as ‘White,’ ‘Black/African American,’ or ‘Other (American Indian, Alaskan Native, Multiracial, or Other) based on reports provided by adult participants and child caregivers.

### DNA extraction and quality analyses

To minimize the impact of DNA extraction procedures, DNA was extracted from all tissues using the Gentra Puregene DNA Extraction Kit according to factory guidelines (Qiagen). This kit has been used to extract DNA from whole blood, PBMCs, saliva, buccal cells, and DBS [50]. Extracted DNA was stored at −80°C in Qiagen DNA Hydration Solution.

Prior to assay for TL, DNA was assessed for integrity, purity, and quantity. DNA integrity and purity were quantified using indicators of DNA degradation from the TapeStation 2200 Bioanalyzer (Agilent) and absorbance ratios from the NanoDrop 2000 spectrophotometer (Thermo Fisher Scientific). DNA concentration was quantified in 3 ways: (a) the NanoDrop spectrophotometer was used to quantify total nucleic acids, (b) the Agilent TapeStation and (c) Quant-iT PicoGreen (Invitrogen) to determine double-stranded DNA concentrations. DNA concentrations as determined by Quant-iT Picogreen were used to standardize the number of telomeres being assessed in each sample. Quality DNA metrics are summarized in **Table 2**.

**Table 2.** Summary of DNA integrity, purity, and quantity metrics.

| Metric | Source | Interpretation |
| --- | --- | --- |
| DNA Integrity Number | TapeStation | Increased DNA degradation as values decrease from 10.0 |
| % Unfragmented DNA | TapeStation | %DNA with length greater than 3,000 bp |
| % Highly Fragmented DNA | TapeStation | %DNA with length between 250 bp – 3,000 bp |
| % Severely Fragmented DNA | TapeStation | %DNA with length less than 250 bp |
| A260/230 ratio | Nanodrop Spectrometer | Increased organic contamination as values deviate (±) from 2.00 |
| A260/280 ratio | Nanodrop Spectrometer | Increased protein contamination as values deviate (±) from 1.80 |
| NanoDrop DNA Concentration | Nanodrop Spectrometer | Concentration of total nucleic acids in ng/μL |
| PicoGreen DNA Concentration | Quant-iT PicoGreen | Concentration of double-stranded DNA in ng/μL |
| TapeStation DNA Concentration | TapeStation | Concentration of double-stranded DNA in ng/μL |

### Assessment of telomere length via qPCR and aTL calculation

TL measurements were generated using the quantitative polymerase chain reaction (qPCR) on DNA extracted from PBMCs, buffy coat, DBS, buccal cells, and saliva. TL in an absolute unit of kilobase pairs (aTL) was measured following a qPCR method originally developed by O’Callaghan and Fenech [51] and adapted by the Shalev Lab [52] using a Rotor-Gene Q thermocycler connected to an uninterruptible power source (CyberPower), which has been shown to decrease variability in TL measured via qPCR [53]. Each qPCR assay consisted of two runs, one quantifying telomere content (T), and a second run quantifying genome copy number (S) using the single copy gene *IFNB1*. The two runs (T & S) were always performed on the same day using the same DNA dilution, which was stored at 4°C between runs (∼2.5 hours).

Estimates of kb telomeric DNA and genome copy number were calculated based on the alignment of each sample with a standard curve. Estimates for the no template control were subtracted from estimates of the analytical samples prior to calculating aTL values. The average kb telomeric DNA estimates and genome copy number estimates across triplicate measurements were used to calculate aTL values: aTL = (Estimated kb Telomeric DNA) / (Estimated Genome Copy Number×92).

To control for inter-assay variability, 5 control samples were assessed on each T run and each S run. The average inter-assay CV for control sample aTL estimates was 8.95%. A pseudo-random selection of 88 samples balanced across tissues (except buccal) was reassessed for explicit purposes of calculating the interclass correlation coefficient (ICC), an indicator of measurement reliability. The ICC across 44 samples rerun for reproducibility was 0.772 (0.728 when a ‘Tissue’ factor was included). The ICC for 44 re-extracted samples was 0.826, which decreased to 0.784 when a ‘Tissue’ factor was added to the model. Full details on qPCR assays for aTL, including reaction mix composition and sequences for primers and standards, are summarized in **S1 Table** in accordance with guidelines recommended by the Telomere Research Network [54].

### Statistical analyses

Statistical analyses were performed using R Studio V2022.07.2 (R 4.1.1). We assessed all continuous variables for skewness and kurtosis. aTL was approximately normal alongside DIN, % unfragmented, % highly fragmented, % severely fragmented, and A260/230 (|skew| < 1; |kurtosis| < 3). However, A260/280 and all three extracted DNA concentrations violated assumptions of normality. Outlier values for each continuous variable were winsorized, where outliers were defined as values outside the range of (Q1-1.5IQR) to (Q3+1.5IQR) across the sample stratified by cohort and tissue, where Q1 and Q3 are lower and upper quartiles respectively, and IQR is the interquartile ratio. Outlier values were winsorized to the boundary values of this range. Winsorizing data points based on the IQR is more appropriate for variables with skewed distributions, in comparison to winsorizing based on standard deviations away from the mean. 295/5891 (5.0%) data points were winsorized across the study (**S2 Table**, see **S1 Fig** for variable distributions before and after winsorization). Results using raw and winsorized data were not statistically different.

To assess biological variation in aTL, we performed a linear mixed effect model [R package nlme; 55] predicting all aTL values with fixed effects of age, sex (female vs. male), tissue (buccal, saliva, DBS, buffy coat, PBMC), race (white, black, other), and an age by tissue interaction, with an additional random effect of individual ID. We included an age by tissue interaction to assess whether tissues differ in chronological age-related changes in aTL [5]. Post-hoc analyses were performed using the *emmeans* package [56]. Using the *correlation* package [57], we also assessed partial Spearman’s correlations of aTL among tissue types within individuals, which accounted for variation in age and sex.

Similar to analysis of aTL values, we performed separate linear mixed effect models predicting each quality DNA metric with fixed effects of age, sex, tissue, race, and an age by tissue interaction, with a random effect of individual ID. We also assessed partial Spearman’s correlations among metrics indicative of DNA integrity (DIN and % fragmentation indices), purity (A260/280, A260/230), and quantity (extracted DNA concentration measured by NanoDrop, PicoGreen, and TapeStation). Partial Spearman’s correlations accounted for age and sex of participants.

We next explored whether DNA metrics of integrity, purity, and quantity predicted aTL, using a two-prong approach. First, we performed partial Spearman’s correlations between aTL and each DNA metric, accounting for age and sex. Second, we performed model comparisons to ask whether certain DNA metrics improved model fits of tissue-specific aTL. We evaluated support for competing candidate models predicting aTL. For each tissue type, we used the *dredge* function [58] to create model sets from the global model (below), in which all models for a given tissue included the same subset of data. Each model could include any combination of age, sex, race, DIN, % unfragmented, highly fragmented, or severely fragmented DNA, A260/280, A260/230, and each of three DNA concentrations, but variables with a correlation above 0.40 were not allowed to coexist in a single candidate model. The number of terms (excluding the intercept) in a single candidate model was limited to approximately 1 term per 10 observations. In addition, TapeStation metrics (DNA integrity and concentration) were not included in candidate models for buffy coat to enhance statistical power because buffy coat was only measured in the child cohort and only 23 children had TapeStation data. We used the Akaike information criterion corrected for small sample sizes (AICc) for model comparisons [59] and present ΔAIC (AIC_i_–AIC_best_ model) and AIC weights (weight of evidence for model) for the top model set, which included models with ΔAIC ≤ 2. Then, we performed conditional model averaging of top model sets.

For each set of models, ANOVA tables are presented in the main text, and coefficient tables are included in the supplemental material. Potential inflation in type I error of multiple statistical testing was controlled separately for each part of analyses using the Benjamini-Hochberg method. P values of statistical significance after controlling for false discovery rate (FDR) at <0.01 were indicated using asterisks in each table or figure that involves statistical testing.

## Results

### Biological variation in aTL

aTL significantly shortened with chronological age (F_1,191_ = 99.15, p<0.001), the magnitude of which varied by tissue type (F_4,557_=15.65, p<0.001, **Fig 1A**, **Table 3A**, **S3 Table**). In particular, post hoc analyses showed significant age-related decreases from 8 to 70 years in aTL for buccal, DBS, and PBMC (buccal: β = −0.12, 95% CI=[-0.15, −0.10]; DBS: β = −0.12, [-0.15, −0.10]; PBMC: β = −0.12, [-0.16, −0.07]), but not for saliva (age 8 to 70 years) or buffy coat (age 8 to 15 years) (saliva: β = −0.02, [-0.05, 0.01]; buffy coat: β = −0.05, [-0.40, 0.31]). Tissues also significantly differed in aTL values (F_4,557_ = 131.89, p < 0.001, **Fig 1B**, **S4 Table**). After adjustment for multiple comparisons, saliva and buccal aTL were significantly shorter than all other tissue types *except* for children buffy coat aTL, which was not significantly different from all other tissues. aTL values of all blood-based tissues (i.e., DBS, buffy coat, and PBMCs) were not statistically different. aTL did not vary by sex (F_1,191_=2.46, p=0.12, **S2 Fig**) or race (F_2,191_ = 1.54, p = 0.22) across all tissue types.

**Fig 1.**
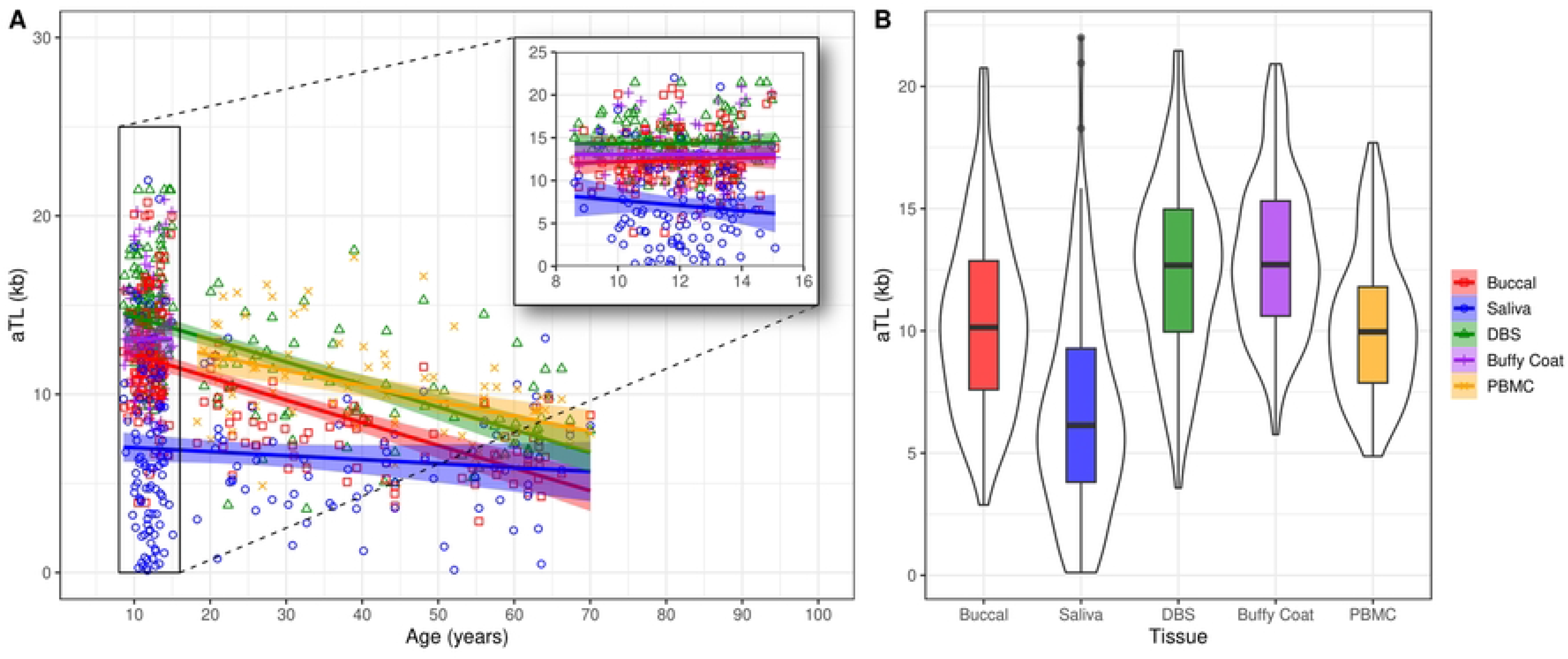
Biological variation in aTL with chronological age (A) and tissue type (B) for individuals ranging from 8 to 70 years old. Note that buffy coat and PBMC are exclusive to child and adult cohorts, respectively.

**Table 3.** Linear mixed effects models predicting aTL and metrics of DNA integrity, purity, and quantity with tissue type and sample demographics. P-values were adjusted for multiple comparisons using the Benjamini-Hochberg method. Asterisks indicate significant p-values after controlling false discovery rate (FDR) at < 0.01. Primary outcomes of interest were analyzed in different models indicated by different panels A-J.

| (A) aTL |  |  |  | (F) A260/280 |  |  |  |
| --- | --- | --- | --- | --- | --- | --- | --- |
| Predictors | df | F | p | Predictor | df | F | p |
| (Intercept) | 1, 557 | 3965.51 | <0.001* | (Intercept) | 1, 577 | 340349.02 | <0.001* |
| Age | 1, 191 | 99.15 | <0.001* | Age | 1, 191 | 15.70 | <0.001* |
| Sex | 1, 191 | 2.46 | 0.119 | Sex | 1, 191 | 2.04 | 0.155 |
| Tissue | 4, 557 | 131.89 | <0.001* | Tissue | 4, 577 | 86.36 | <0.001* |
| Race | 2, 191 | 1.54 | 0.216 | Race | 2, 191 | 0.87 | 0.419 |
| Age x Tissue | 4, 557 | 15.65 | <0.001* | Age x Tissue | 4, 577 | 20.24 | <0.001* |
| (B) DNA Integrity Number (DIN) |  |  |  | (G) A260/230 |  |  |  |
| (Intercept) | 1, 280 | 29497.55 | <0.001* | (Intercept) | 1, 577 | 4568.29 | <0.001* |
| Age | 1, 94 | 22.33 | <0.001* | Age | 1, 191 | 46.77 | <0.001* |
| Sex | 1, 94 | 0.91 | 0.343 | Sex | 1, 191 | 1.49 | 0.224 |
| Tissue | 4, 280 | 212.95 | <0.001* | Tissue | 4, 577 | 45.03 | <0.001* |
| Race | 2, 94 | 3.95 | 0.023 | Race | 2, 191 | 0.35 | 0.707 |
| Age x Tissue | 4, 280 | 1.32 | 0.264 | Age x Tissue | 4, 577 | 3.17 | 0.014 |
| (C) %Unfragmented DNA (> 3000 bp) |  |  |  | (H) Nanodrop Concentration (ng/μL) |  |  |  |
| (Intercept) | 1, 288 | 15664.94 | <0.001* | (Intercept) | 1, 577 | 740.35 | <0.001* |
| Age | 1, 94 | 14.20 | <0.001* | Age | 1, 191 | 0.65 | 0.421 |
| Sex | 1, 94 | 1.42 | 0.237 | Sex | 1, 191 | 0.06 | 0.801 |
| Tissue | 4, 288 | 173.18 | <0.001* | Tissue | 4, 577 | 113.41 | <0.001* |
| Race | 2, 94 | 2.12 | 0.125 | Race | 2, 191 | 0.55 | 0.578 |
| Age x Tissue | 4, 288 | 2.86 | 0.024 | Age x Tissue | 4, 577 | 3.22 | 0.013 |
| (D) %Highly Fragmented DNA (250–3000 bp) |  |  |  | (I) PicoGreen Concentration (ng/μL) |  |  |  |
| (Intercept) | 1, 288 | 997.72 | <0.001* | (Intercept) | 1, 577 | 684.23 | <0.001* |
| Age | 1, 94 | 5.59 | 0.02 | Age | 1, 191 | 0.00 | 0.996 |
| Sex | 1, 94 | 1.33 | 0.252 | Sex | 1, 191 | 0.16 | 0.694 |
| Tissue | 4, 288 | 133.65 | <0.001* | Tissue | 4, 577 | 188.55 | <0.001* |
| Race | 2, 94 | 1.46 | 0.238 | Race | 2, 191 | 0.19 | 0.825 |
| Age x Tissue | 4, 288 | 4.65 | 0.001* | Age x Tissue | 4, 577 | 1.40 | 0.231 |
| (E) %Severely Fragmented DNA (< 250 bp) |  |  |  | (J) TapeStation Concentration (ng/μL) |  |  |  |
| (Intercept) | 1, 288 | 817.11 | <0.001* | (Intercept) | 1, 288 | 322.02 | <0.001* |
| Age | 1, 94 | 2.52 | 0.116 | Age | 1, 94 | 1.25 | 0.267 |
| Sex | 1, 94 | 0.63 | 0.43 | Sex | 1, 94 | 2.15 | 0.146 |
| Tissue | 4, 288 | 79.57 | <0.001* | Tissue | 4, 288 | 105.48 | <0.001* |
| Race | 2, 94 | 1.27 | 0.286 | Race | 2, 94 | 0.39 | 0.678 |
| Age x Tissue | 4, 288 | 4.26 | 0.002* | Age x Tissue | 4, 288 | 2.86 | 0.024 |

aTL values were significantly correlated between all tissue pairs *except* PBMC-buccal (⍴ = 0.21) and PBMC-saliva (⍴ = 0.18), as well as correlations between buffy coat and saliva (⍴ = 0.22, **Fig 2**). Partial Spearman’s ⍴ values for all the pairs ranged from 0.18 (PBMC-saliva) to 0.51 (PBMC-DBS). Several of the stronger correlations occurred between related tissues, e.g., DBS-buffy coat and DBS-PBMC in the child and adult cohorts, respectively. Excepting buccal-saliva correlations, which were significant in adults (⍴ = 0.41), but not children (⍴ = 0.26), tissue pair correlations did not significantly differ if separated by cohort (see **S3 Fig**).

**Fig 2.**
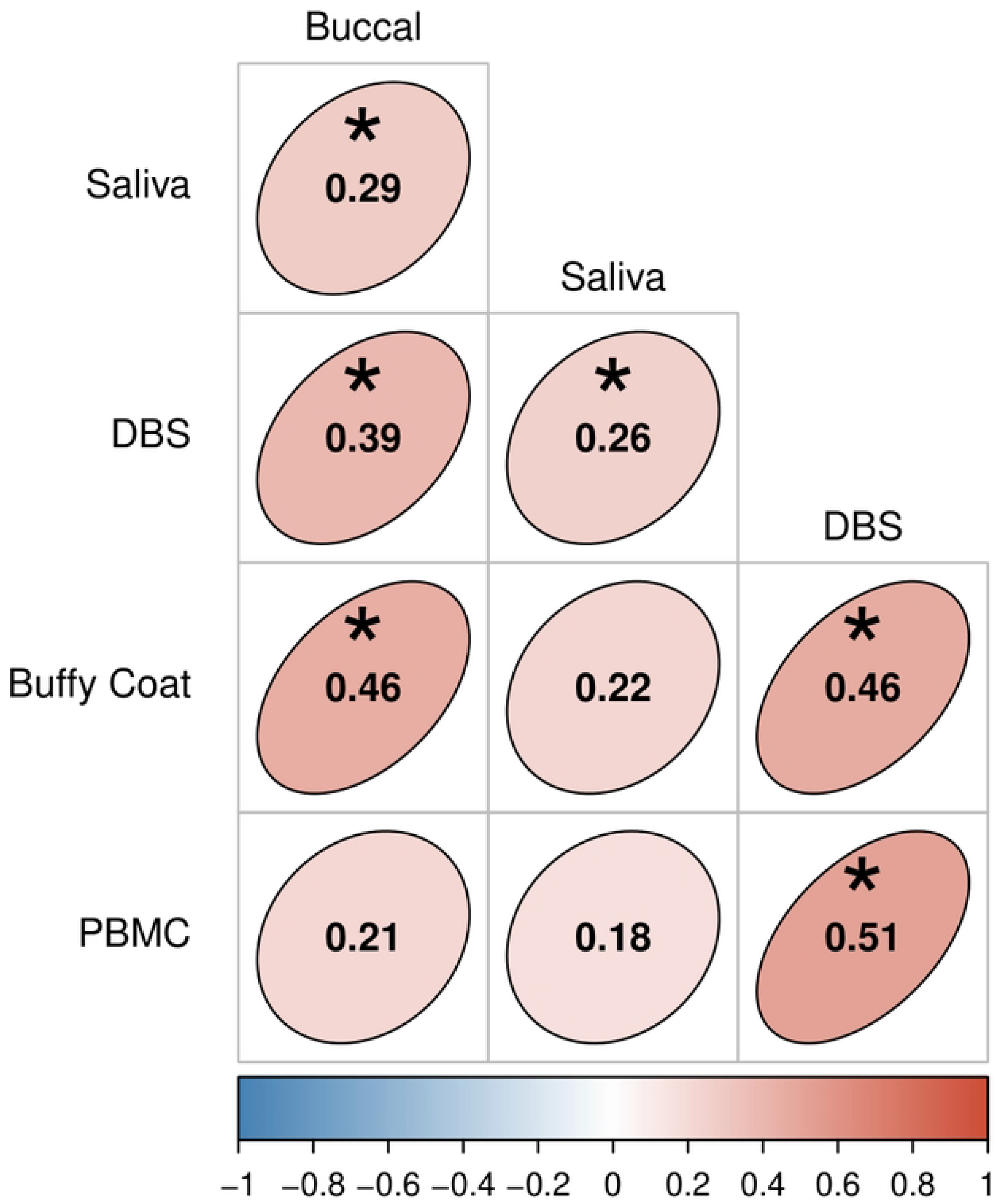
Partial Spearman’s correlations of aTL among tissue types, which account for age and sex. Ellipse shape and color denotes the strength and direction of correlations. Asterisks indicate significant p-values after adjusting for multiple comparisons using the Benjamini-Hochberg method and controlling false discovery rate (FDR) at < 0.01.

### Biological variation in DNA metrics of integrity, purity, and quantity

All results describing variation in DNA metrics can be found in **Fig 3**, **Tables 3-4**, and **S4-S5 Tables**. DIN values significantly varied by tissue type (F_4,280_ = 212.95, p < 0.001, **Fig 3A-D**) and are mirrored by patterns of % DNA fragmentation (unfragmented: F_4,288_ = 173.18, p < 0.001, highly fragmented: F_4,288_ = 133.65, p < 0.001; severely fragmented: F_4,288_ = 79.57, p < 0.001). Notably, buccal DIN values were lowest among all tissues (DIN_mean_ = 5.6). Interestingly, DIN and % unfragmented DNA appear higher in samples from older participants (DIN: F_1,94_ = 22.33, p < 0.001; unfragmented: F_1,94_ = 14.20, p < 0.001). A260/280 values also varied by tissue type (F_4,577_ = 86.36, p < 0.001, **Fig 3E**), where DBS had significantly lower A260/280 values than all other tissue types. A260/280 values were lower in older participants (F_1,191_ = 15.70, p < 0.001), although this varied by tissue (F_4,577_ = 20.24, p < 0.0001). A260/230 values also significantly differed by tissue type (F_4,577_ = 48.163, p < 0.001, **Fig 3F**); PBMCs had significantly higher A260/230 than all other tissues except for buffy coat. A260/230 values were significantly lower in older participants (F_1,191_ = 46.77, p < 0.001). All DNA concentration types significantly varied among the majority of tissue pairs (NanoDrop: F_4,577_ = 113.41, p < 0.001; PicoGreen: F_4,577_ = 188.55, p < 0.001; TapeStation: F_4,577_ = 105.48, p < 0.001; **Fig 3H-J**), with DBS/saliva and buffy coat/PBMC exhibiting the lowest and highest concentrations, respectively. DNA metrics did not vary by sex or race.

**Fig 3.**
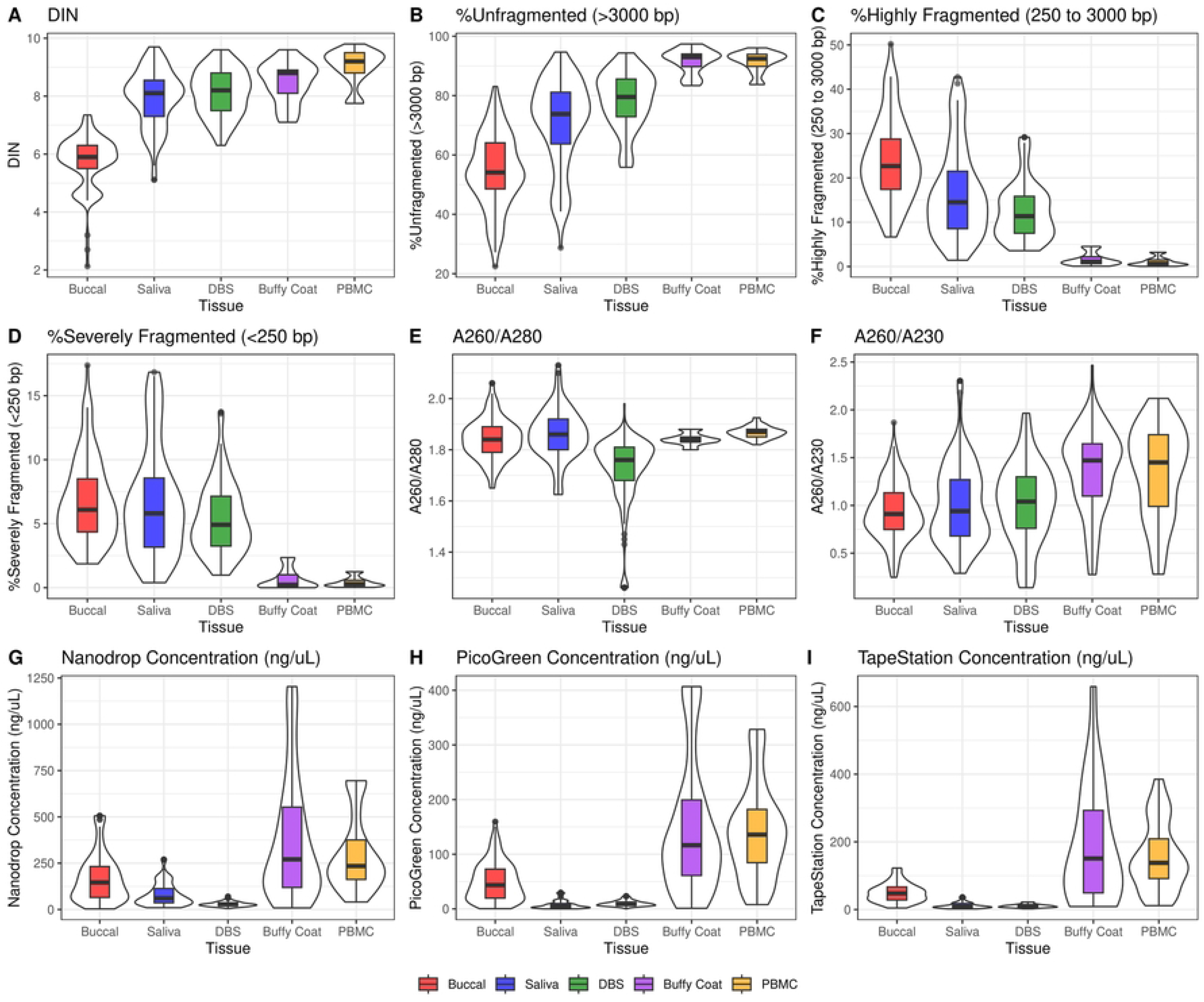
Variation in metrics of DNA integrity (A-D), purity (E-F), and quantity (G-I) across tissue types.

**Table 4.** Tissue-level averages of aTL and metrics of DNA integrity, purity, and quantity, split by child and adult cohorts. Values are presented as tissue/cohort averages with standard error in parentheses.

|  | Buccal |  | Saliva |  | DBS |  | Buffy Coat | PBMC |
| --- | --- | --- | --- | --- | --- | --- | --- | --- |
| Variable | Child | Adult | Child | Adult | Child | Adult | Child | Adult |
| aTL (kb) | 12.39<br>(3.25) | 7.45<br>(1.99) | 7.10<br>(4.86) | 6.05<br>(2.90) | 14.38<br>(2.86) | 9.75<br>(3.06) | 13.08<br>(3.19) | 10.27<br>(2.96) |
| DIN | 5.31<br>(1.31) | 5.89<br>(0.71) | 7.43<br>(1.16) | 8.08<br>(0.87) | 7.41<br>(0.59) | 8.33<br>(0.74) | 8.47<br>(0.71) | 9.02<br>(0.59) |
| % Unfragmented DNA<br>(> 3000 bp) | 50.25<br>(12.68) | 55.73<br>(11.98) | 64.5<br>(15.57) | 72.83<br>(13.99) | 75.22<br>(7.87) | 79.61<br>(9.87) | 91.58<br>(4.16) | 91.41<br>(3.47) |
| % Highly Fragmented DNA<br>(250 – 3000 bp) | 27.44<br>(9.05) | 22.59<br>(8.96) | 20.26<br>(11.19) | 15.48<br>(9.61) | 11.56<br>(4.23) | 12.71<br>(6.91) | 1.67<br>(1.48) | 1.10<br>(0.99) |
| % Severely Fragmented DNA<br>(<250 bp) | 8.01<br>(3.38) | 6.36<br>(3.13) | 7.64<br>(4.54) | 6.32<br>(4.12) | 4.61<br>(1.68) | 5.62<br>(2.98) | 0.67<br>(0.82) | 0.43<br>(0.38) |
| A260/A280 | 1.87<br>(0.08) | 1.80<br>(0.06) | 1.84<br>(0.09) | 1.90<br>(0.10) | 1.77<br>(0.07) | 1.65<br>(0.18) | 1.84<br>(0.02) | 1.87<br>(0.02) |
| A260/A230 | 1.05<br>(0.28) | 0.75<br>(0.21) | 1.13<br>(0.45) | 0.84<br>(0.29) | 1.19<br>(0.33) | 0.76<br>(0.41) | 1.36<br>(0.43) | 1.35<br>(0.48) |
| Nanodrop Concentration<br>(ng/μL) | 164.67<br>(147.06) | 178.38<br>(99.04) | 64.11<br>(51.35) | 107.58<br>(67.23) | 28.88<br>(10.21) | 29.49<br>(16.25) | 381.64<br>(335.82) | 297.27<br>(198.29) |
| PicoGreen Concentration<br>(ng/μL) | 47.61<br>(39.15) | 54.44<br>(29.23) | 5.15<br>(4.75) | 11.28<br>(9.14) | 10.04<br>(4.46) | 9.98<br>(5.64) | 149.97<br>(120.79) | 141.24<br>(85.26) |
| TapeStation Concentration<br>(ng/μL) | 48.84<br>(27.84) | 50.98<br>(28.79) | 9.76<br>(5.77) | 12.65<br>(9.24) | 11.91<br>(3.78) | 10.27<br>(5.14) | 188.96<br>(173.32) | 156.22<br>(100.64) |

Many metrics of DNA integrity, purity, and quantity were moderately to strongly correlated. Full results are shown in **Fig 4**, **S4 Fig**, and **S5 Table**, but we highlight key patterns here. First, DIN values were strongly correlated with DNA fragmentation indices for all tissue types, with the exception of buffy coat, for which we had limited power. Absolute ⍴ values ranged from 0.19 to 0.95, where high DIN values were characterized by a higher proportion of unfragmented DNA. In addition, all extracted DNA concentrations were significantly positively correlated for all tissues except buffy coat (0.37 < ⍴ < 0.94; ⍴_mean_ = 0.70). Interestingly, higher extracted DNA concentrations were linked to higher DIN values, particularly for DNA concentrations measured via TapeStation. For NanoDrop and PicoGreen concentrations, correlations are strongest for saliva and DBS (0.16 < ⍴ < 0.81; ⍴_mean_ = 0.60). Concentration of extracted DNA was also positively associated with A260/230 in all tissues except DBS; however, A260/280 exhibited inconsistent associations with DNA quantity, with absolute values of ⍴ ranging from 0.03 to 0.56. DIN metrics were inconsistently related to A260/280 and A260/230.

**Fig 4.**
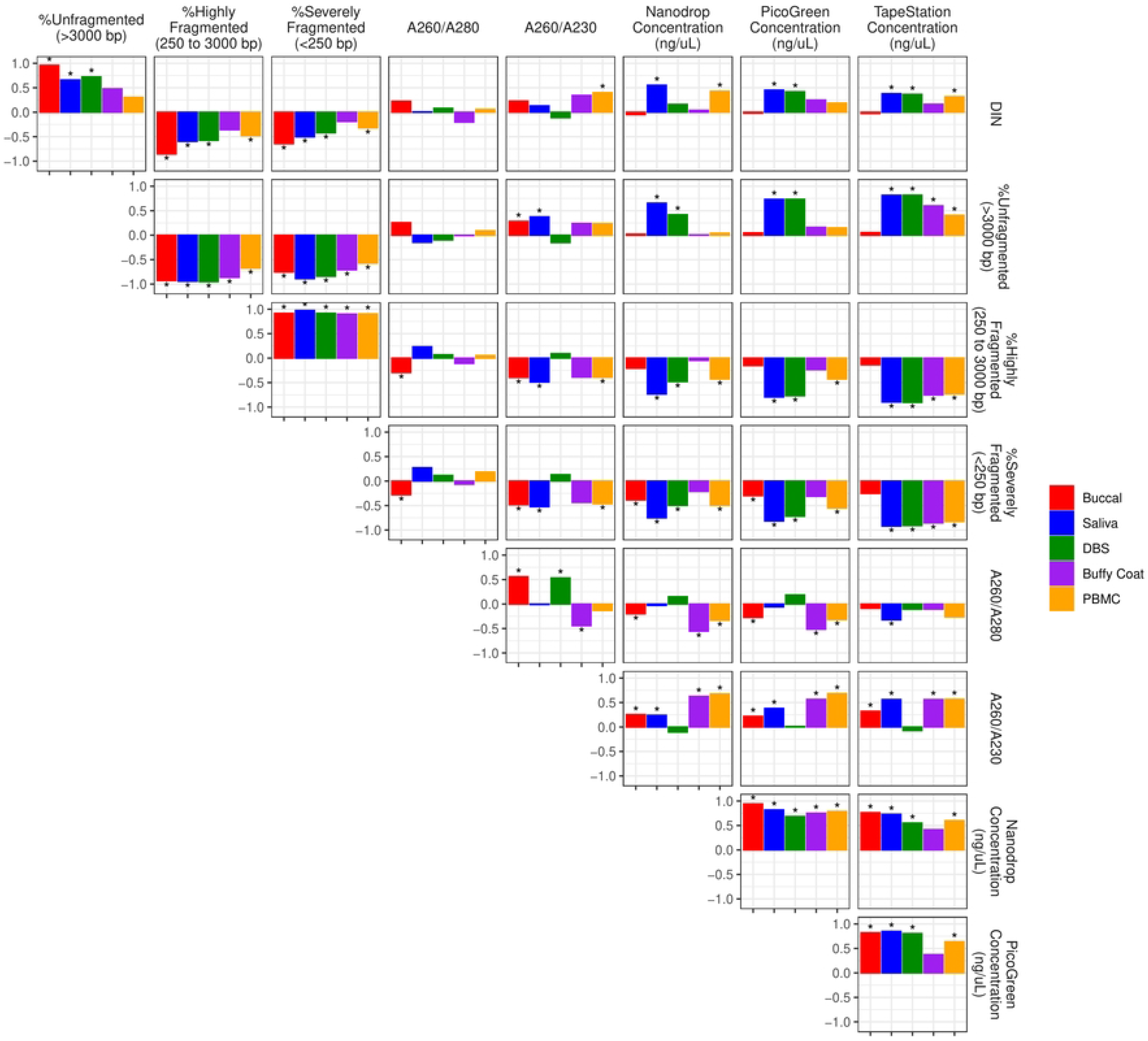
Partial Spearman’s correlations among DNA metrics for each tissue type, after accounting for age and sex of participants. Spearman’s *⍴* values range from −1 to 1 on the y-axis. Asterisks indicate significant p-values after adjusting for multiple comparisons using the Benjamini-Hochberg method and controlling false discovery rate (FDR) at < 0.01.

### Covariation between aTL and metrics of DNA integrity, purity, and quantity

Partial Spearman’s correlations showed that aTL is significantly correlated with DNA integrity values in some tissues (**Fig 5**, **S5 Fig**, **S6 Table**). While aTL is overall weakly and inconsistently correlated with DIN and DIN-related metrics, higher DIN or low % fragmentation is significantly associated with longer aTL in saliva and PBMCs. In addition, aTL is significantly and positively correlated with all three DNA concentrations across most tissues, ranging from 0.02 < ⍴ < 0.62, particularly so in saliva, buccal, and buffy coat. Correlations between aTL and A260/280 were overall weak, and A260/230 was only significantly associated with aTL in buccal and buffy coat. Overall, longer aTL is associated with lower % DNA fragmentation, higher extracted DNA concentrations, and higher A260/230. We also note that correlations between DNA metrics and aTL appear particularly strong for saliva.

**Fig 5.**
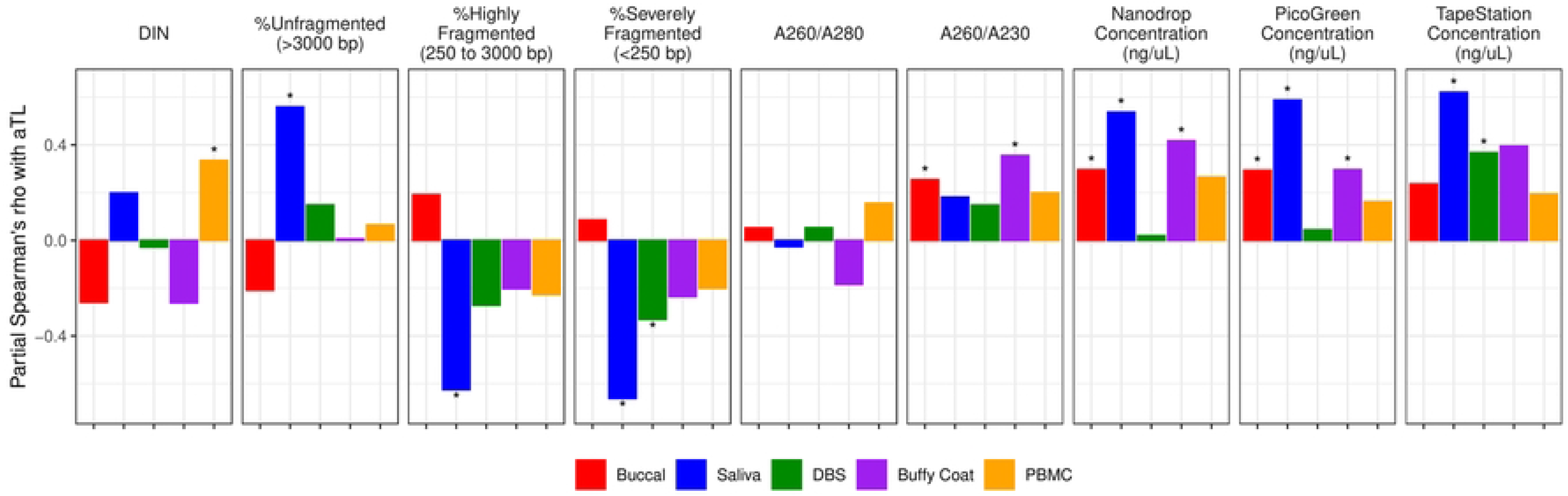
Partial Spearman’s correlations between aTL and metrics of DNA integrity, purity, and quantity, adjusted for age and sex and split by tissue type. Spearman’s *⍴* values range from −1 to 1 on the y-axis. Asterisks indicate significant p-values after adjusting for multiple comparisons using the Benjamini-Hochberg method and controlling false discovery rate (FDR) at < 0.01.

Results for model comparisons can be found in **Table 5** and **S7 Table**. Among candidate models predicting aTL in buccal, the top-ranked model set included DIN, % highly fragmented DNA, and A260/230 as significant predictors of aTL. TapeStation/PicoGreen DNA concentrations were also included in the top-ranked model set but did not significantly predict buccal aTL. The top-ranked model set in saliva only included % severely degraded DNA and A260/280, for which only the former had high variable importance and significantly predicted aTL. The top-ranked model set in DBS included DIN, A260/280, A260/230, and TapeStation DNA concentration, and all variables but DIN significantly predicted aTL after conditional averaging. The top-ranked model set predicting buffy coat aTL only included NanoDrop DNA concentration as a significant predictor (TapeStation metrics were not included in models for buffy coat). The top-ranked model set in PBMC included DIN, % unfragmented and severely fragmented DNA, A260/280, and TapeStation and NanoDrop concentrations, but only DIN and TapeStation concentration predicted PBMC aTL. Across all tissues, ΔAIC values for null intercept-only models were ≥ 17.00 and for null age-only models, were ≥ 7.85 (**Table S7**), suggesting that inclusion of DNA metrics significantly improved model fits of aTL beyond that of chronological age alone. However, there were no consistent variables across tissues in the top model sets.

**Table 5.** Conditional model-averaged coefficients for the top models sets (ΔAIC ≤ 2) investigating the relative importance among DNA metrics in improving model fit of aTL values, split by tissue type. For each DNA metric in the top model set, we also provide variable importance (VIMP), or the sum of model weights across all top models that contain each DNA metric, standardized by the sum of model weights of the top model set. A VIMP value equal to 1 means that variable was present in all models in the top model set. For race, B/O refer to estimates of aTL for Blacks and Other relative to Whites.

|  | Buccal |  | Saliva |  | DBS |  | Buffy Coat |  | PBMC |  |
| --- | --- | --- | --- | --- | --- | --- | --- | --- | --- | --- |
| | $\beta$ (SE) | VIMP | $\beta$ (SE) | VIMP | $\beta$ (SE) | VIMP | $\beta$ (SE) | VIMP | $\beta$ (SE) | VIMP |
| Age | -0.05 (0.01)* | 1.00 | -0.05 (0.02)* | 1.00 | -0.08 (0.02)* | 1.00 | -0.09 (0.18) | 0.18 | -0.08 (0.20)* | 1.00 |
| Sex | -0.49 (0.40) | 0.35 | -0.49 (0.60) | 0.19 |  |  | -0.61 (0.55) | 0.25 | -1.42 (0.61) | 1.00 |
| Race | B 1.81 (1.01) | 0.21 | 3.86 (1.69) | 0.78 | 3.17 (1.45) | 0.55 |  |  |  |  |
|  | O 0.13 (0.62) |  | -0.14 (0.99) |  | -0.45 (0.88) |  |  |  |  |  |
| DIN | -1.16 (.23)* | 0.44 |  |  | -0.54 (0.39) | 0.53 |  |  | 1.39 (0.54) | 0.67 |
| %Unfragmented<br>(> 3000 bp) |  |  |  |  |  |  |  |  | 0.12 (0.10) | 0.15 |
| %Highly Fragmented<br>(250–3000 bp) | 0.12 (0.02)* | 0.56 |  |  |  |  |  |  |  |  |
| %Severely<br>Fragmented<br>(<250 bp) |  |  | -0.63 (0.09)* | 1.00 |  |  |  |  | -0.74 (0.89) | 0.15 |
| A260/280 |  |  | -2.99 (3.62) | 0.19 | 6.51 (1.98)* | 0.12 |  |  | 20.83 (15.39) | 0.44 |
| A260/230 | 6.17 (0.92)* | 1.00 |  |  | 2.60 (0.75)* | 0.88 |  |  |  |  |
| TapeStation | 0.01 (0.01) | 0.37 |  |  | 0.26 (0.07)* | 1.00 |  |  | 0.003 (0.003) | 0.15 |
| PicoGreen | 0.008 (0.01) | 0.06 |  |  |  |  |  |  |  |  |
| Nanodrop |  |  |  |  |  |  | 0.004 (0.001)* | 1.00 | 0.005 (0.002)* | 0.32 |

\*p < 0.01 after FDR

## Discussion

We assessed tissue variation in aTL in a cross-sectional dataset of 8- to 70-year-old individuals. To our knowledge, this is one of a few studies to compare TL between a selection of invasively and non-invasively sampled tissues in a cohort that includes both children and adults. aTL significantly shortened with chronological age for all tissues except saliva and buffy coat, the latter of which had a restricted age range (i.e., 8 to 15 years). aTL varied by tissue, particularly between blood and non-blood tissues. Despite this variation, aTL was correlated across most tissue pairs. We also observed variation in metrics of DNA integrity, purity, and quantity and explored whether controlling for such variation improved predictions of aTL. Many metrics were correlated: higher extracted DNA concentration was associated with higher DIN and more acceptable A260/230 values. DNA metrics varied by tissue, and blood-based tissues (especially PBMC and buffy coat) had higher integrity and quantity DNA. Cross-tissue variation in DNA qualities may help drive variation in aTL, and we provided evidence that longer aTL is linked to higher DIN, DNA concentrations, and to some extent, A260/230 values. Model comparisons suggest that incorporation of DNA metrics significantly improves predictions of aTL, although important metrics vary by tissue. These results highlight potential considerations for tissue selection in future population-based studies of TL and the value of incorporating quality DNA metrics as control variables to improve TL prediction.

Tissues significantly differed in aTL values and age-related changes in aTL. In particular, non-invasively sampled tissues (buccal cells and saliva) had shorter aTL than blood-based tissues [similar to 35]. This does contrast with other work in which saliva TL is longer than blood [60, 61]; however, methodological differences may drive this discrepancy. Tissue type often maps onto variation in TL [34–37] and is likely due to tissue-specific cell composition and turnover rates, stem ‘cellness’, and TL maintenance [28, 30, 33]. Similar TL regulation among related tissues may explain why aTL of blood-based tissues were similar, and such physiology may also influence rates of TL attrition. Here, all tissues *except* saliva and buffy coat shortened with age: aTL of buccal, DBS, and PBMC decreased by ∼120 bp/year, but only by 18 and 48 bp/year for saliva and buffy coat, respectively. 120 bp/year is higher than previous estimates, i.e., well below 100 bp/year for most tissues [34, 62]. Null associations between age and aTL buffy coat could be explained by a narrow age range within the child cohort (8-15 years).

While aTL decreased with chronological age for most tissue types, it was not significantly linked with other external validity metrics, including sex and race. Previous work often reveals longer TL in females than males [48, 63], although this pattern varies across vertebrates [47]. Here, sex differences may be masked by the relatively larger variation in aTL among tissue types. In addition, TL is often found to be longer in individuals self-identifying as non-Hispanic Black relative to non-Hispanic White [2, 49, but see 64], an effect that we cannot fully test due to the limited racial/ethnic diversity of participants in this study.

Complementing the rapidly-growing number of TL studies in epidemiology is additional research on the consequences of variation in TL methodology on measurement validity and research outcomes [24–26], including sample collection, storage, extraction, and TL measurement assay. Yet, whether and how sample-specific metrics of DNA quality influence TL is unexplored. DNA degradation and amount are used to predict genotyping success [65] and has become particularly relevant for degraded forensic samples [66]. Similarly, poorer-quality DNA may interfere with telomere assay precision and/or yield inaccurately short TL values. Here, assessing variation in quality DNA metrics has revealed several patterns.

First, tissues differed in DNA integrity, purity, and quantity. Results show that blood-based tissues (buffy coat and PBMCs) had higher quality DNA, namely higher and less variable DNA integrity, less variable A260/280, more acceptable A260/230, and higher extracted DNA concentrations. On the other hand, buccal cells and DBS had the lowest DIN and A260/280 values, respectively. Few other studies have compared DNA metrics by tissue, but Lucena-Aguilar [41] showed that DNA purity and integrity were lower in formalin-fixed paraffin-embedded tissues compared to frozen tissues and saliva. In addition, Hansen et al. [40] showed that DNA quality was highest in blood, and surprisingly saliva, when compared to DNA from buccal cells. Interestingly, DIN was higher and A260/280 was lower in older individuals, although the former could be an artifact driven by high PBMC (adult-only tissue) DIN. Age differences may also stem from age-related changes in cell composition or amount and ease of tissue collection [67].

Second, many metrics of DNA integrity, purity, and quantity were significantly correlated. As expected, high DIN values were associated with increased percentages of unfragmented DNA, and DNA concentration was correlated across all three quantification methods (i.e., NanoDrop, PicoGreen, and TapeStation). Interestingly, high extracted DNA concentrations for the majority of tissue types were associated with high DNA integrity and A260/230, the latter of which has been shown in human saliva [41]. This may be expected if we assume that samples with high extracted DNA concentrations come from tissues with higher cellular density, as exemplified by the higher DNA concentrations of buffy coat and PBMCs vs non-blood tissues, and relative to DBS cards, which were collected from whole blood and thereby included a large proportion of non-nucleated red blood cells. In this case, samples with increased cellular density (and higher DNA concentration) may degrade less during storage and extraction and be less susceptible to organic or protein contamination. Given that DNA integrity may influence telomere assays, it may therefore be important to minimize variation in and correlations among DNA metrics by standardizing sample inputs during extraction by volume and cell counts.

Next, we assessed whether variation in quality metrics of DNA improved models of aTL. Interestingly, longer aTL was associated with lower % DNA fragmentation, higher DNA concentrations, and more acceptable (or closer to 2.0) A260/230. That the extracted DNA concentration predicts aTL *despite* a standardized amount of DNA being put into TL reactions suggests that controlling for or reducing variation in extracted DNA concentration could be vital to decreasing noise in aTL outputs. Interestingly, saliva aTL appears consistently and strongly associated with DNA metrics (i.e., DIN, A260/230, DNA concentration), and so incorporating these metrics may be vital in certain tissue types. In fact, model comparisons show that incorporation of DNA metrics into aTL models significantly improved model fit, as age-only null models had much greater ΔAIC values than models with age and DNA metrics. However, across tissues, there were no quality metrics of DNA that appeared more often in top-ranked sets, i.e., most DNA metrics appeared in 2-3 tissues’ top-ranked model sets. Tissues exhibiting a low-quality ‘tail’ for a specific DNA metric were more likely to have that DNA metric appear as predictive of aTL for that tissue. For example, buccal and DBS have low-DIN and low-A260/280 ‘tails’, respectively, and here, their aTLs are significantly related to those metrics. Future studies should continue to assess the importance of quality metrics of DNA to improve models of TL.

We acknowledge certain limitations of this study. First, tissue types collected from the child and adult cohorts were unbalanced. The child cohort did not have PBMCs isolated from whole blood, while the adult cohort did not have buffy coat. This restricted the age range of the dataset when evaluating cross-tissue and cross-age variations of aTL and DNA metrics, which may explain the non-significant shortening of TL with age observed in buffy coat. Second, TapeStation metrics were not measured for all child samples, which limited the power to examine their associations with age and aTL, especially in buffy coat, a child-only tissue. Additionally, we did not control for several factors that may induce variation in aTL, including blood cell proportions for blood-based tissues [68] and factors like exposures and lifestyles that are linked to TL dynamics in previous work [69, 70].

How might this information inform future population-based studies of TL? As shown in limited previous work [41], blood-based samples exhibited the highest quality DNA and therefore, may be preferred for reliable measurement of TL. Buffy coat and PBMCs exhibited high DNA integrity and more acceptable A260/280 and A260/230 values compared to less invasive tissues like buccal and saliva, which appear to exhibit more variable and lower quality DNA metrics. DBS, as a minimally invasive tissue, had similar aTL values to PBMC and buffy coat, and can be an alternative to blood-based samples, especially in pediatric populations. Saliva in particular had lower DNA integrity and aTL values that were strongly influenced by metrics of DNA quality and did not significantly decrease with age despite being measured in both the child and adult cohorts. That previous work supports saliva as an acceptable alternative to blood [40, 41] conflicts with our results and suggests the need for additional tissue comparisons of DNA quality metrics. However, not all new or ongoing studies can rely on blood-based tissues. In this case, our results show that quantifying sample-specific metrics of DNA quality for use in model predictions of TL can improve model fits of the data, thereby strengthening the signal of exogeneous predictors of TL and the utility of TL as a proxy for health-related outcomes. Alternative to controlling for variation in DNA metrics, standardizing DNA extractions to yield consistent concentrations could also minimize methodological impacts on TL measures. We encourage further study of variation in quality metrics of DNA across tissues and how it may mediate variation in TL, which can help inform how to select tissues and/or control for differences in DNA quality in future population-based telomere studies.

## Acknowledgements

We thank the children and caregivers for their participation in the study, Child Health Study staff, all nurses at the CRC and the adult participants in this study.

## Supporting Information

**S1 Table. Telomere Research Network Reporting Guidelines**

**S2 Table. Number of winsorized data points for each continuous variable, split by cohort and tissue.** Outliers are defined as values outside the range of (Q1-1.5IQR) to (Q3+1.5IQR) for each cohort-tissue subset of data points, where Q1 and Q3 are lower and upper quartiles respectively, and IQR is the interquartile ratio. Outlier values were winsorized to the boundary values of this range. 295/5891 (5.0%) datapoints were winsorized across the study.

**S3 Table**. **Summary of coefficient outputs for models predicting aTL and metrics of DNA integrity, purity, and quantity with tissue type and sample demographics.**

**S4 Table. Contrasts between tissues for each dependent variable, including aTL and metrics of DNA integrity, purity, and quality.** Asterisks indicate significant p-values after adjusting for multiple comparisons using the Benjamini-Hochberg method and controlling false discovery rate (FDR) at < 0.01.

**S5 Table. Partial Spearman’s ⍴ values for correlations between metrics of DNA integrity, purity, and quantity, as shown in Fig 3 in the main text.**

**S6 Table. Partial Spearman’s ⍴ values for correlations between metrics of DNA integrity, purity, and quantity and aTL.**

**S7 Table. Top model sets (ΔAICc ≤ 2) for models predicting aTL with age, sex, race, and metrics of DNA integrity, purity, and quantity**, in which no predictors were correlated above ⍴ = 0.4. k = number of parameters in each candidate model, including the intercept; wi = Akaike model weight. Intercept-only and age-only null models are highlighted in gray for each tissue.

**S1 Fig. Histogram distributions of continuous variables of interest, before and after winsorization (gray and blue distributions, respectively).** A datapoint was winsorized if it fell outside the range of (Q1-1.5IQR) to (Q3+1.5IQR) for its respective cohort-tissue distribution of data points, where Q1 and Q3 are lower and upper quartiles respectively, and the IQR is the interquartile ratio. Outlier values were winsorized to the boundary values of this range. 375/6673 (5.6%) datapoints were winsorized across the study.

**S2 Fig**. **Biological variation in aTL with tissue type and sex for individuals ranging from 8 to 70 years old.** Buffy coat and PBMC are exclusive to child and adult cohorts, respectively.

**S3 Fig**. **Partial Spearman’s correlations of aTL among tissue types, accounting for age and sex and split by child and adult cohorts.** Ellipse shape and color denotes the strength and direction of correlations. Significant correlations (p < 0.05) are indicated by an asterisk. Buffy coat and PBMC are exclusive to the child or adult cohort, respectively.

**S4 Fig**. **Partial Spearman’s correlations among metrics of DNA integrity, purity, and quantity, split by cohort and tissue type, after accounting for age and sex of participants.** Y-axis *p* values range from −1 to 1, and significant correlations (p < 0.05) are indicated by an asterisk.

**S5 Fig**. **Partial Spearman’s correlations between aTL and metrics of DNA integrity, purity, and quantity, adjusted for age and sex and split by tissue and cohort.**

## Notes

### Competing Interest Statement

The authors have declared that no competing interests exist.

